# TRUmiCount: Correctly counting absolute numbers of molecules using unique molecular identifiers

**DOI:** 10.1101/217778

**Authors:** Florian G. Pflug, Arndt von Haeseler

**Affiliations:** Max F. Perutz Laboratories (MFPL), Center for Integrative Bioinformatics Vienna (CIBIV), University of Vienna, Medical University of Vienna, Vienna, Austria; Bioinformatics and Computational Biology, Faculty of Computer Science, University of Vienna, Vienna, Austria

## Abstract

Counting DNA or RNA molecules using *next-generation sequencing* (NGS) suffers from amplification biases. Counting *unique molecular identifiers* (UMIs) instead of reads is still prone to over-estimation due to amplification and sequencing artifacts and under-estimation due to lost molecules. We present an algorithm that corrects for these errors, based on a mechanistic model of the PCR and sequencing process whose parameters have an immediate physical interpretation and are easily estimated. We demonstrate that our algorithm outputs essentially unbiased counts with substantially improved accuracy.

---

Experimental methods like RNA-seq, ChIP-Seq and many others depend on NGS to measure the abundance of DNA or RNA molecules in a sample. The PCR amplification step necessary before sequencing often amplifies different molecules with different efficiencies, thereby biasing the measured abundances. This problem can be alleviated by ensuring that all molecules are distinguishable before amplification by some combination of factors comprising a *unique molecular identifier* (UMI)^1^, which usually includes a distinct molecular barcode ligated to each molecule before amplification (figure 1A, 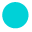, 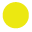, 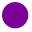, 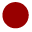, 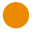). After amplification and sequencing, instead of counting reads, reads are grouped by UMI, and each distinct UMI is taken to reflect a distinct molecule in the original sample (figure 1A). But while the number of distinct UMIs may be a better proxy for the molecule count, it is still biased, for two reasons:

a. Molecules that are amplified with low efficiency will have fewer copies made, hence fewer reads per UMI, and thus a higher chance of being left entirely unsequenced (figure 1A, 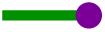).
b. Sequencing errors, PCR chimeras, and index miss-assignment^2^ in multiplexed sequencing runs can produce *phantom UMIs* which do not correspond to any molecule in the original sample (figure 1A, 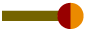).

**Figure 1:**
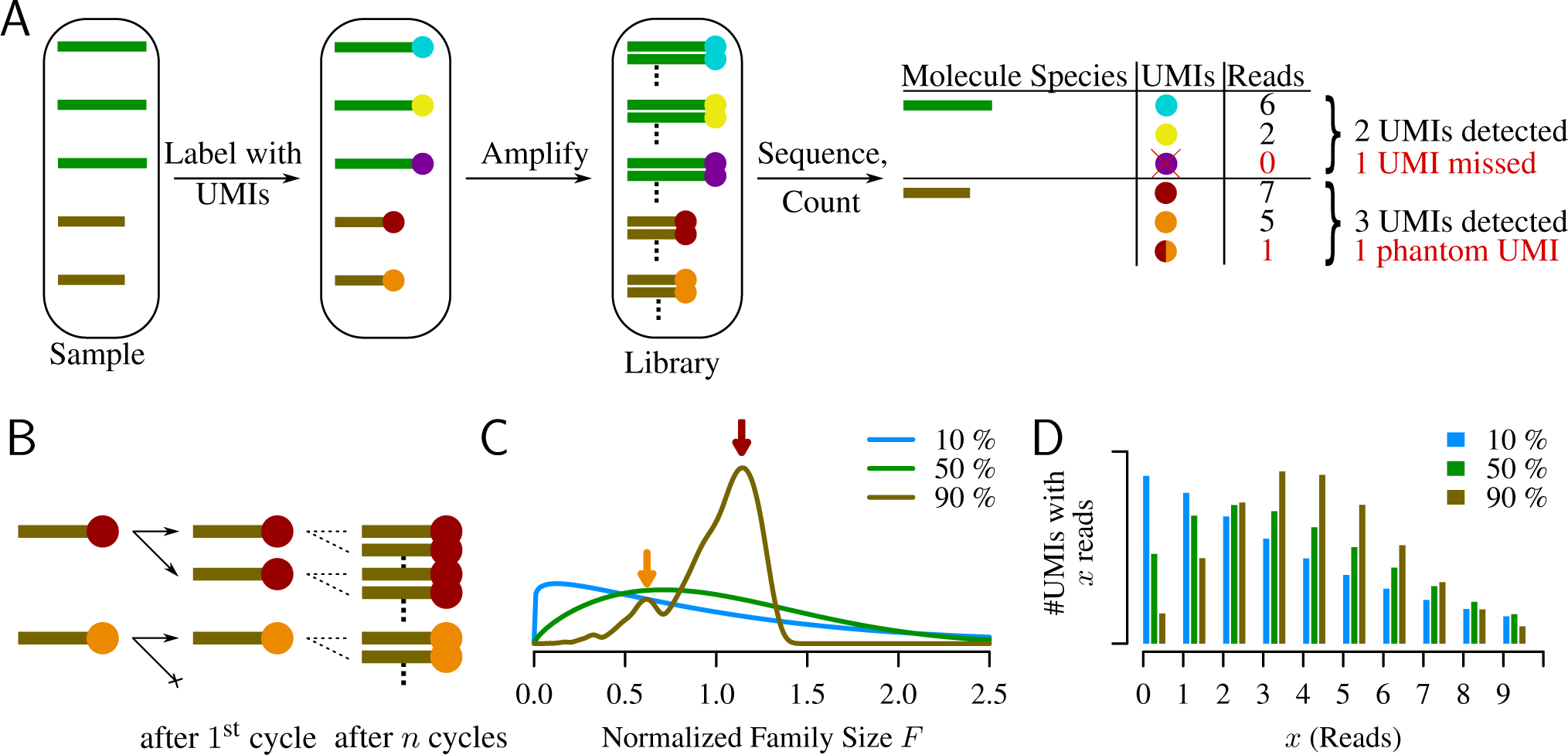
**A**. The relevant steps of library preparation when the UMI method is used. The sample initially contains 3 copies of molecule 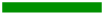 and 2 copies of 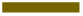, which are made unique by labeling with UMIs (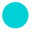, 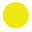, 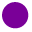, 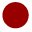, 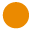). Each of those molecules is expanded into a molecular family during amplification, and a random selection of molecules from these families are sequenced. Counting unique UMIs then counts unique molecules, unless UMIs have read-count zero (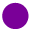) or phantom UMIs are produced (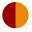). **B**. PCR as a Galton-Watson branching process. Molecule 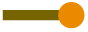 failed to be copied during the 1^st^ cycle and the final family size is thus reduced compared to 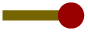. **C**. Normalized family size distribution for efficiency 10%, 50% and 90%. The arrows mark the most likely normalized family sizes for the two molecules from (B), assuming a reaction efficiency of 90%, and taking their distinct fates during the 1^st^ cycle into account. **D**. Distribution of reads per UMI for efficiency 10%, 50% and 90% assuming *D* = 4 Reads per UMI on average.

Chimeric PCR products are typically produced during later reaction cycles, and can therefore be expected to have smaller copy numbers and hence a lower readcount than non-chimeric PCR products. Index miss-assignment and sequencing errors typically happen randomly, and are unlikely to produce a larger number of reads showing the same phantom UMI. For these reasons, phantom UMIs can be expected to have a markedly lower read count than most *true UMIs*, i.e. UMIs of actual molecules in the original sample.

We present the three-step bias-correction and phantom-removal algorithm *TRUmiCount* that exploits this difference in expected read counts:

1. We first filter out phantom UMIs by removing any UMI whose read count lies below a suitable chosen *error-correction threshold* (*T*).
2. We then estimate the loss (*f*), i.e. the fraction of molecules that were not sequenced at all, or whose UMIs were removed by the error-correction threshold. This estimate is computed using a stochastic model of the amplification and sequencing process whose parameters are the PCR efficiency (*E*), and the sequencing depth (*D*), expressed as the average number of reads per UMI in the initial sample. From the observed distribution of reads per UMI, we estimate both (raw) gene-specific as well as library-wide values for these parameters, and compute corresponding estimates of the loss.
3. Finally, we add the estimated number of lost UMIs back to the the observed number of true UMIs (those UMIs with ≥ threshold reads) to find the total number of molecules in the original sample. Since the loss can vary between genes, to yield unbiased counts, the correction must be based on gene-specific loss estimates. Because the raw gene-specific estimates are noisy for genes with only few observed true UMIs, we employ a James-Stein-type^3^ *shrinkage estimator*, adjusting the raw gene-specific parameter and loss estimates towards the library-wide ones (thus *shrinking* their difference). We choose the amount of shrinkage based on each estimate’s precision, in such a way that the expected overall error is minimized^4^.

We start the construction of our model of the amplification and sequencing process with a model of PCR amplification introduced by Krawczak *et al.*^5^. Here, each molecule is assumed to be duplicated in each cycle with probability *E*, also called the reaction’s *efficiency* (figure 1B). While this model has been extended by Weiss & von Haeseler^6^ to include the possibility of substitution errors during amplification, exhaustively modeling *all* possible sources of phantom UMIs seems futile. We therefore pursue a different approach, and model only the error-free case, trusting the error-correction threshold to remove phantoms. Over multiple cycles, each molecule is thus assumed to be expanded into a *molecular family* of identical copies. Since each UMI is initially represented by a single molecule, random successes or failures to copy during early cycles lead to a variation in the final family sizes, even between identical (expect for their molecular barcode) molecules. As the family size of each initial molecule grows, the proportion of successful copy operations approaches the efficiency *E*, therefore reducing the amount of noise added by each additional cycle. The total number of cycles thus has little influence on the final family size distribution, and is therefore not a parameter of our model. The final distribution does, however, depend strongly on the reaction efficiency, with fluctuations in family size decreasing as the efficiency grows towards 100%.

For efficiencies close to 100%, most molecular families are thus of about average size, except for those (approximately 100 − *E* percent) families for which the first copy failed. These are about half the average size, and form a distinct secondary peak in the family size distribution (figure 1C, brown curve). We emphasize that due to this, even at efficiencies close to 100%, the distribution still shows considerable dispersion, meaning that even at high efficiencies stochastic PCR effects are not negligible. At lower efficiencies, the family sizes vary even more wildly, as extreme family sizes (on both ends of the scale) become more likely (figure 1C, blue and green curves).

To complete our model of amplification and sequencing, we combine the stochastic PCR model outlined above with a model of sequencing as random Poissonian sampling^7^. The variability of per-UMI read counts thus has two sources – the variability of molecular family sizes and the Poissonian sampling introduced by sequencing. While the latter is reduced by increasing the sequencing depth, the former is independent of the sequencing depth but is reduced by increasing the reaction efficiency. For all reasonable error-correction thresholds *T* the predicted fraction of true UMIs filtered out by the error-correction step thus grows with diminishing efficiency *E*.

We validated this model using two published RNA-seq datasets. Kivioja *et al.^1^* labeled and sequenced transcripts in *D. melanogaster* S2 cells using 10bp random molecular barcodes from the 5’ end. Shiroguchi *et al.^8^* labeled and sequenced transcript fragments in *E. coli* cells on both ends, using (on each end) one of 145 molecular barcodes carefully selected to have large pairwise edit distances. The Y-shaped sequencing adapters used in the *E. coli* experiment were designed such that each strand of a labeled double-stranded cDNA molecule produces a related but distinguishable molecular family.

To see whether our algorithm offers an advantage over existing UMI errorcorrection strategies, we pre-filtered the observed UMIs in each of the two replicates of these datasets using the following existing algorithms: We first merged UMIs likely to be erroneously sequenced versions of the same molecule, using the algorithm proposed by Smith *et al.*^9^. For the *E. coli* experiment we also removed UMIs for which the complementary UMI corresponding to the second strand of the same initial template molecule was not detected, as proposed by Shiroguchi *et al.*^8^.

To this pre-filtered set of UMIs we then applied our algorithm. Above the errorcorrection threshold (figure 2A, black bars), the observed library-wide distribution of reads per UMI closely follows the model prediction, and the *E. coli* data even shows traces of the secondary peak that represents molecules not duplicated in the first reaction cycle. We thus conclude that our model captures the main stochastic behavior of the amplification and sequencing processes, and accurately models the read-count distribution of true UMIs.

**Figure 2:**
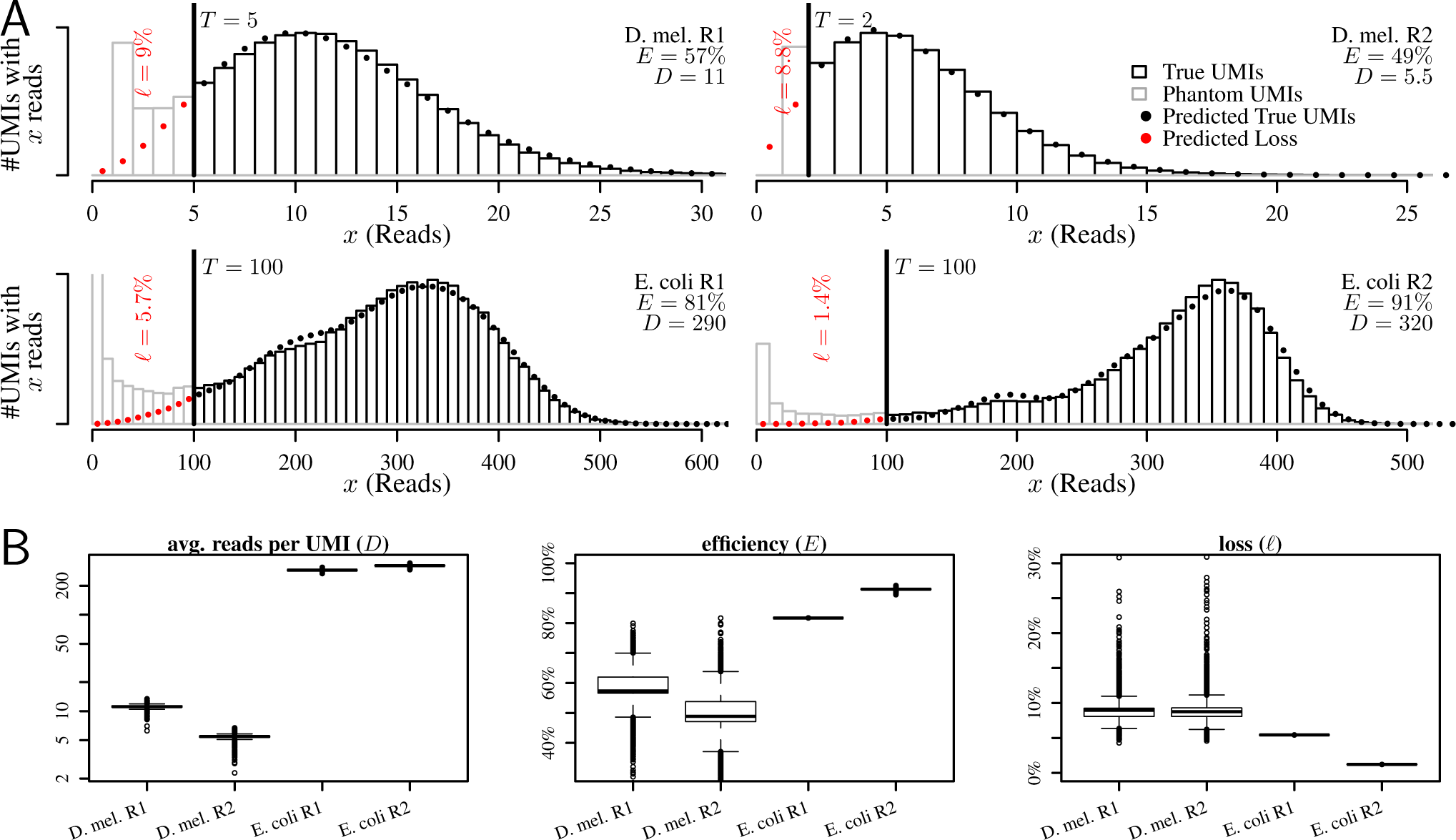
**A**. Observed and predicted library-wide distribution of reads per UMI and parameter and loss estimates. Filtered UMIs (grey bars, left of threshold *T*) are over-abundant and thus assumed to contain both phantom and true UMIs (red dots). UMIs surviving the filter (black bars) closely follow the predicted distribution (black dots) and are assumed to be true UMIs. **B**. Variability of the (shrunken) model parameters and resulting loss between genes. Includes parameter for 7481 detected genes in *D. mel.* R1, 8001 genes in R2, 2380 genes in *E. coli* R1 and 2308 genes in R2.

The UMIs removed by our filter, i.e. those with fewer reads than the errorcorrection threshold demands, (figure 2A, grey bars) are over-abundant compared to our prediction. This over-abundance increases further as per-UMI read counts drop, indicating the existence of a group of UMIs with significantly reduced molecular family sizes. While we may expect some systematic variation of family sizes between true UMIs (on top of the stochastic variations that our PCR model predicts), we would expect these to be gradual and not form distinct groups. We conclude that the UMIs contributing to the observed over-abundance are indeed phantoms that are rightly removed by our algorithm. We note that none of these phantoms were removed by either the UMI merging algorithm of Smith *et al.*^9^, or (for the *E. coli* data) by filtering UMIs for which the complementary UMI (representing the second strand of the template molecule) was not detected.

The gene-specific (shrunken) estimates for amplification efficiency, average reads per UMI, and loss, that our algorithm produces, vary between genes to different degrees (figure 2B). We observe the smallest amount of variation for the average number of reads per UMI (figure 2B, left) – the estimates of this parameter are virtually identical for a large majority of genes, and differs only for a few outliers.

The estimated amplification efficiencies on the other hand can vary substantially between genes, for the two *D. melanogaster* replicates from ≈ 50% up to ≈ 70% (figure 2B, middle). Considering that in this experiment only the 3’ ends of transcripts were sequenced, and all fragments contributing to a gene hence share a similar sequence composition, this is not unexpected. These differences in efficiency cause the loss to vary heavily between genes as well (figure 2B, right), from ≈ 5% in the best case to ≈ 30% in the worst case. Without gene-specific loss corrections, abundance comparisons between genes will thus suffer from systematic biases against certain genes of up to ≈ 30% − 5% ≈ 25%.

In contrast, fragments from all parts of the transcript were sequenced in the *E. coli* experiments, and together with the high sequencing depth (≈ 300 reads per UMI), we now expect little variations of efficiency, and small and highly uniform losses across genes. Our efficiency and loss estimates reflects this (figure 2B, middle and right), and as the lack of outliers shows, they do so even for genes with only few UMIs. Yet for these genes, the raw (unshrunken) gene-specific estimates are noisy (figure S1), proving that shrinking the raw estimates successfully reduces the noise to acceptable levels.

To further verify the accuracy of the corrected transcript counts computed by our algorithm, we conducted a simulation study. We use the (loss-corrected) estimated total transcript abundances of *D. melanogaster* replicate 1, rounded to 10, 30, 100, 300, 1000, 3000 or 10000 molecules as the true transcript abundances. We then simulated amplification and sequencing of these transcripts, using for each gene the previously estimated gene-specific efficiency and average number of reads per UMI (figure 2B). To the resulting list of UMIs and their read-counts for each gene we applied our algorithm to recover the true transcript abundances (threshold *T* = 5 as before), and determined for each gene the relative error of the recovered abundances compared to the simulation input.

Figure 3 shows these relative errors (a) if no correction is done (b) if the correction is based soley on the raw gene-specific loss estimates (i.e. no shrinkage) and (c) for the complete algorithm as presented (i.e. using shrunken loss estimates). The uncorrected counts systematically under-estimate the true transcript counts, in 50% of the cases by at least ≈ 10%, independent of the true number of transcripts per gene. And even at high transcript abundances, the relative error still varies *between* genes, biasing not only absolute transcript quantification, but also relative comparisons between different genes. The counts corrected using raw gene-specific estimates are unbiased and virtually error-free for strongly expressed genes, but exhibit a large amount of additional noise for weakly expressed genes. The complete presented algorithm using shrunken loss estimates successfully controls the amount of added noise, and shows no additional noise for weakly expressed genes, while still being unbiased and virtually error-free for more strongly expressed genes.

**Figure 3:**
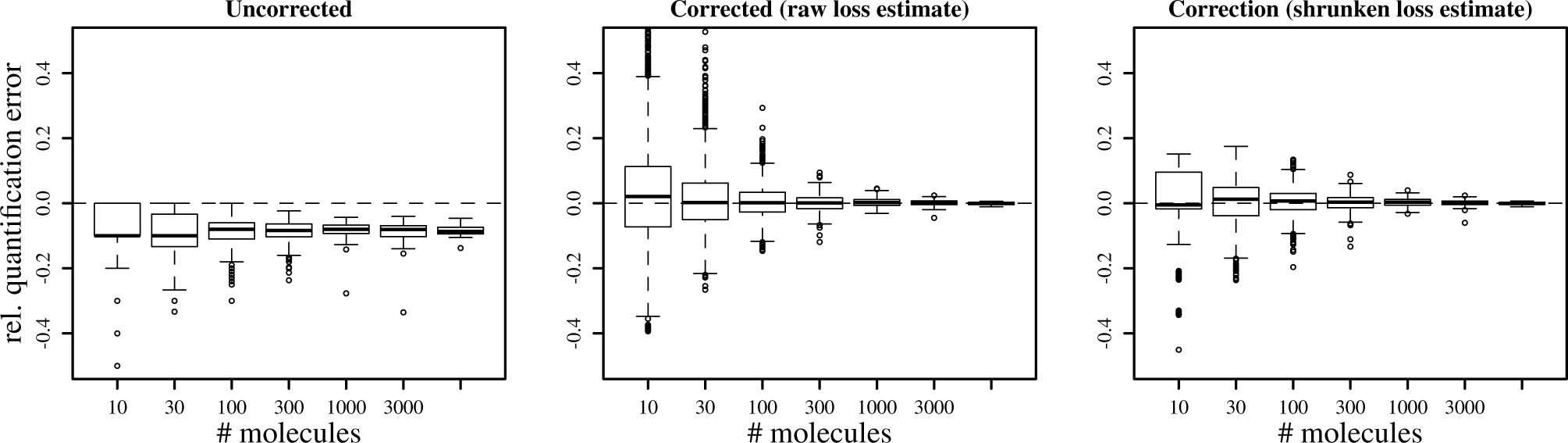
Relative error of estimated total number of UMIs depending on the true number of UMIs. Left panel uses the observed number of UMIs without any correction. Middle panel uses the raw gene-specific loss estimates to correct for lost UMIs. Right panel uses the shrunken gene-specific estimates to correct for losses.

Thus, the *TRUmiCount* algorithm we presented successfully removes the biases inherent in raw UMI counts, and produces unbiased and low-noise measurements of transcript abundance, allowing for unbiased comparisons between different genes, exons, and other genomic features. It does so even in the presence of various types of phantom UMIs and varying amplification efficiencies, both between samples and along the genome. Compared to other error-correction techniques, it is not restricted to particular types of phantom UMIs, or to a special Y-shaped design of the sequencing adapters.

Our model of the amplification and sequencing process is mechanistic, and its two parameters have an immediate physical interpretation. They can both be determined from the experimental data without the need for either guesses or separate calibration experiments. The *TRUmiCount* algorithm thus does not require any changes to library preparation over the basic UMI method. By inspecting the estimated parameters – in particular the amplification efficiency, the amplification reaction itself can be studied. For example, by estimating model parameters separately for sequenced fragments of different lengths, the drop of reaction efficiency with increasing fragment lengths can be quantified (figure S3).

The *TRUmiCount* algorithm can thus help to increase the accuracy of many quantitative applications of NGS, and by removing biases from comparisons between genes can aid in the quantitative unravelling of complex gene interaction networks. To make our method as easily accessible as possible to a wide range of researchers, we provide two readily usable implementations of our algorithm. Our R package *gwpcR* enables a flexible integration into existing R-based data analysis workflows. In addition, we offer the command-line tool *TRUmiCount* which is designed to work in conjunction with the *UMI-Tools* of Smith *et al.*^9^. Together they provide a complete analysis pipeline which produces unbiased transcript counts from the raw reads produced by an UMI-based RNA-Seq experiment (http://tuc:tuc@www.cibiv.at/~pflug_/trumicount).

## Acknowledgements

We would like to thank all members of the *Center for Integrative Bioinformatics Vienna* (CIBIV), in particular Luis Paulin-Paz, Celine Prakash and Philipp Rescheneder for their valuable feedback throughout all stages of this project, and Olga Chernomor for sharing her insights into shrinkage estimators.

## Author contributions

Both authors contributed equally to this work.

## METHODS

### A stochastic model of PCR

We follow the *single-stranded* model of Krawczak *et al.*^5^ and view PCR as a stochastic process that during each cycle duplicates each molecule independently with a particular probability *E*, called the reaction’s *efficiency*. For simplicity, we further assume that a molecule is copied perfectly or not at all, i.e. that neither partial copies nor copies with a slightly different base-pair sequence are produced, that no molecules are destroyed or lost, and that the efficiency *E* stays constant throughout the reaction. Since we use the single-stranded model, *molecule* for us always means a single-stranded piece of DNA, and we do not distinguish between a strand and its reverse complement. For our purposes, a piece of double-stranded DNA thus consists of two indistinguishable molecules.

Before amplification, we assume all molecules in the sample to be distinguishable by some UMI. During amplification, each of those molecules gives rise to a *molecular family* of (indistinguishable) copies. The initial size of such a family (i.e. the number of copies it is comprised of) is 1. During the first cycle, the size increases to 2 if the single initial molecule is copied successfully, i.e. with probability *E*. Continuation of this process, always using all existing molecules as potential templates that are copied with probability *E*, produces a random sequence *M*_0_, *M*_1_, *M*_2_,…of molecular family sizes after the 0^th^, 1^st^, 2^nd^, … cycle. This sequence forms a Galton-Watson branching process^10^, and follows the recursion

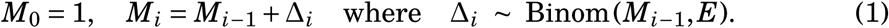

While we are not aware of a way to obtain an explicit formula for the distribution of the family size *M_i_* after *i* cycles, the expected value and variance of *M_i_* can be computed explicitly. According to Harris^11^ (Ch. 1, Eq. 5.3), 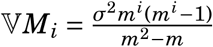 where *m* respectively *σ* are the mean respectively standard deviation of *M*_1_. In our case these are *m* = 1 + *E* respectively *σ*^2^ = *E* ⋅ (1 − *E*) and we find

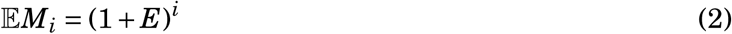

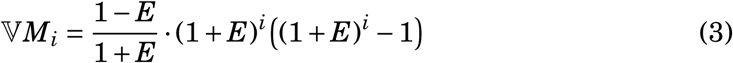

Equation 2 shows the well-known exponential growth of expected family sizes during PCR. But apart from recovering this well-known property of PCR, the Galton-Watson model also predicts the likelihood of *deviations* from this expectation due to random failures of copy operations, and by simulation allows us to find the actual distribution of *M_i_*.

### The normalized family size *F*

Due to the exponential growth of the expectation of *M_i_*, the distribution of *M_i_* depends heavily on the cycle count *i*. That dependency, however, effects mostly only the *scale*, not the *shape* of the distribution of *M_i_*. To see the effects on the shape more clearly, the effects on the scale is removed by replacing *M_i_* with a re-scaled version which has an expected value of one,

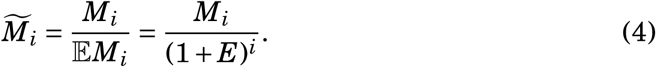

These re-scaled family sizes can be sensibly compared across cycles. We observe that with growing cycle counts, the additional stochasticity introduced by each additional cycle drops rapidly. The re-scaled family size after the first cycle varies by a factor of two depending on whether the (single) copy operation during the first cycle succeeds or fails. Later on there are more templates to copy from, and thus the success or failure to copy any particular molecule averages out, making the behavior of the reaction more deterministic. Finally, *M*̃_i_ ≈ *M*̃_*i*+1_, because the family size *M_i_* increases during each cycle almost exactly by a factor of 1 + *E*, which matches the decrease of the re-scaling factor in *M*̃*_i_*. This informal argument can be turned into a formal proof (see Harris^11^, Ch. 1, Th. 8.1) of the convergence of the re-scaled family size as *i* tends towards *∞*, which allows us to remove the cycle count as a parameter entirely from what we call the *normalized family size*

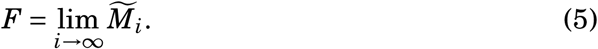

While there is again no explicit formula known for the distribution of the normalized family size *F*, we find its variance from equations 3, 4 and 5 to be

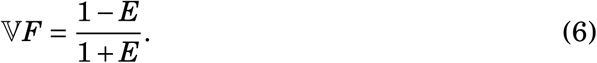

### Computing the distribution of *F*

To find the actual distribution (in terms its density *d*_PCR_(⋅|*E*) with the reaction efficiency *E* as a parameter) of the normalized family size *F* for a particular efficiency *E* we resorted to simulation. We simulated the PCR process for efficiencies from 0.01 to 0.99 (steps of 0.01 up to 0.90, steps of 0.005 up to 0.94, steps of 0.002 up to 0.99). Each time, we simulated 10^9^ independent trajectories, and ran each simulation until the expected family size was 107 molecules (i.e. for *n* = 7/ log_10_(1 + *E*) cycles). At that point the stochasticity further cycles would introduce is negligible and we may thus assume *M*̃*_n_*≈ *M*̃_*n*+1_ ≈ *F*.

For each efficiency *E*, we normalized the simulated raw family sizes using equation 4 to obtain 10^9^ independent samples of *F*. Using kernel density estimation, we then estimated values of the density function *d*_PCR_(*λ | E*) of the normalized family size distribution on a grid of 318 values of *λ* between 0 and 50. The grid points are spaced non-uniformly, being finest (distance 0.0025) around 0 and 1 and getting coarser elsewhere.

This procedure resulted in a 123 *×* 318 matrix of densities, i.e. *d*_PCR_(*λ | E*) evaluated for each combination of one of the 123 simulated efficiencies *E*, and one of the 318 normalized family sizes *λ*. Using this (pre-computed and stored) matrix, the density function *d*_PCR_(*λ | E*) can be evaluated quickly for arbitrary values of *E* and *λ* by two-dimensional polynomial interpolation^12^.

### The Sequencing Process

The normalized family size distribution models the abundance of molecules with a particular UMI. To model the read count of a particular UMI after sequencing (i.e. the number of reads stemming from a particular pre-amplification molecule), we model next-generation sequencing with a poissonian sampling model^7^. This amounts to assuming that (a) each individual copy has the same probability of being sequenced, (b) this probability is small compared to the sequencing depth and (c) there were many (distinguishable) original molecules. We further assume that an UMI is *on average* represented by *D* reads. Then the read count *C* of an UMIs with known normalized molecular family size *F* is Poisson distributed,

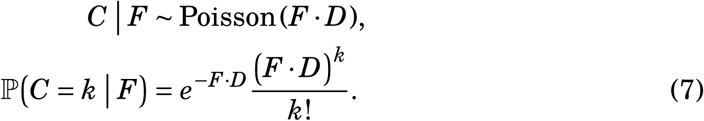

In general, however, the exact family size *F* of any particular UMI is unknown – we only know the *distribution* of *F*. To compute the probability of an UMI having coverage *k*, we average over all possible family sizes, taking their respective probabilities according to the stochastic PCR process into account,

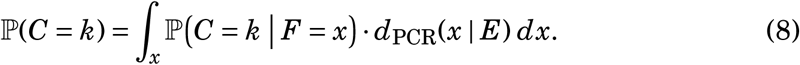

The probabilities ℙ(*C = k*) can be computed efficiently by numerical integration.

Since we impose an error-correction threshold *T* and drop UMIs with fewer than *T* reads, the read-count distribution we actually observe is a *censored* version of *C* where the possible outcomes *C < T* are removed. For the mean and variance of this censored distribution with threshold *T* we write

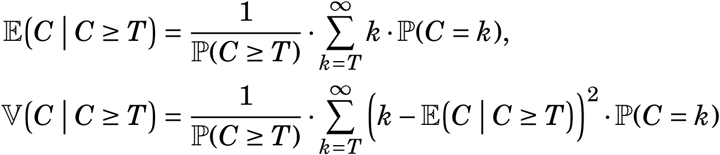

In general, we compute the censored mean and variance numerically. For the uncensored case *T* = 0 (were even unsequenced molecules would somehow be observed) we explicitly find

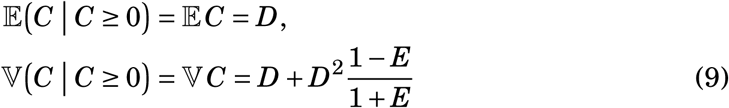

The expected *loss* 𝓁 is the expected fraction of true UMIs that either remain completely unsequenced, or that are removed by the error-correction threshold, meaning

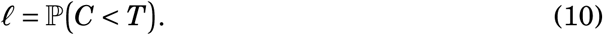

### Estimating parameters and correcting for loss

Given *n*^obs^ experimentally observed UMIs (after applying the error-correction threshold *T* to filter out phantoms) and their read count vector **c** = (*c*_1_,…, *c*_*n*^obs^_), we estimate the reaction efficiency *E* and the mean number of reads per UMI *D*. We use the *method of moments*, i.e. we find *E* and *D* such that the predicted mean equals the sample mean 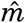 of **c**, and the predicted variance its sample variance 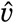. Since we only take observed UMIs with at least *T* reads into account, we must compute the predictions using the censored distribution, i.e. find *E*, *D* such that

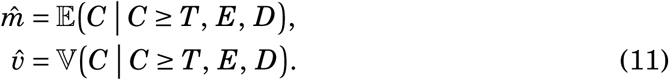

If *T* = 0, i.e. if 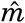 and 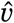 reflect the *uncensored* mean respectively variance, these equations can be solved explicitly by inverting equation 9, which yields the method of moments estimator

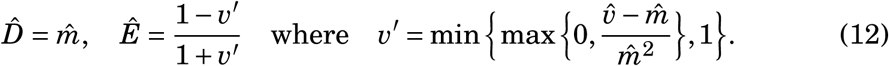

If *T* > 0, we solve the system of equations numerically to find *E* and *D*. With these parameter estimates, we then compute an estimate 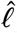 of the loss 𝓁 using equation 10, and use it to correct for the expected number of lost molecules. Assuming that we observed *n*^obs^ UMIs and given 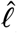, we estimate the total number of molecules in the original sample to have been

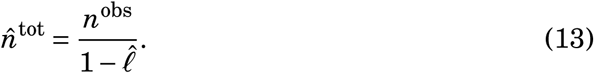

### Gene-specific estimates & corrections

Since the reaction efficiency *E* and depth *D*, and hence also the loss, will usually vary between individual genes (or other genomic features of interest), to correct the observed number of transcripts of some gene *g ∈* 1,…, *K* for the loss, a *gene-specific* loss estimate 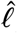*_g_* should be used. In principle, such estimates are found by applying the described estimation procedure to only the UMIs found for transcripts of gene *g*, i.e. by computing a gene-specific mean 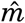*_g_* and variance 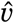*_g_* of the number of reads per UMI, solving equations 11 to find a gene-specific 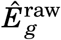 and 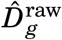, and computing 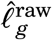 using equation 10. If the number 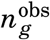 of observed UMIs (i.e. transcripts) stemming from gene *g* is large, a correction based on 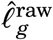 yields an (approximately) unbiased and accurate estimate of the total number of transcripts of that gene. But if 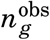 is small, the error of the estimator 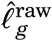 easily exceeds the variability of the true gene-specific value 𝓁_*g*_ between genes. In such cases, correcting using the *library-wide* estimate 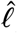^all^ computed from *all* UMIs found in the library will yield an more accurate (although biased) estimate of the total number transcripts of gene *g*.

Somewhat surprisingly, by combining these two flawed estimators of the true gene-specific loss 𝓁*_g_*, we obtain a so-called *shrinkage estimator* 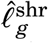 that improves upon both in terms of *mean squared error* (MSE, Carter & Rolph^4^ Eq. 2.4),

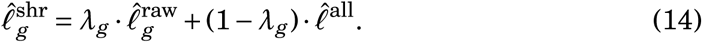

The gene-specific coefficient *λ_g_* determines how much the raw gene-specific estimate is *shrunk* towards the global estimate, and its optimal choice (with respect to the MSE) depends on the variances the two constituent estimators. To determine the optimal *λ_g_* we make the following assumptions about these estimators:

a. the library-wide estimate 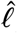^all^ is a good proxy for the true *average* loss taken over all genes 1,…, *K*. This seems reasonable given the size of a typical library, comprising millions of UMIs.
b. the estimation error of the raw gene-specific estimator 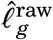 depends only on the number 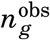 of observed UMIs for gene *g*, and does so in an inversely proportional manner. This is certainly true asymptotically for large numbers of observations, for small numbers it should still provide a reasonable approximation.

We write *s* for the variance of the true loss between genes (i.e. for the mean squared difference of 𝓁*_g_* and 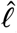^all^), and *u* for the proportionality constant between the error of 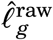 and 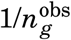. According to Carter & Rolph^4^ (Eq. 2.4 ff.) the optimal choice for *λ_g_* is then

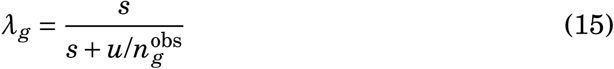

To compute the gene-specific shrinkage estimators 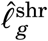, it remains to find constants *u* and *s*. Towards that end, we observe that the expected squared deviation of the raw gene-specific loss estimate 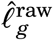 from its average 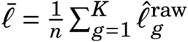 is the total variance of 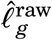, which is comprised of the between-gene variance *s* and the estimator variance 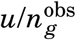, or in other words

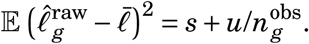

This allows us to estimate *s* and *u* using *least squares regression*, i.e. by minimizing

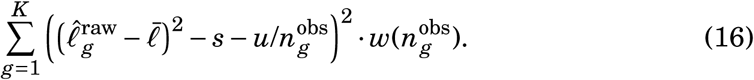

Without weighting (i.e. for *w*(*n*) = 1), the considerable drop in magnitude of 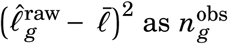 increases would allow genes with small number of observations to yield an unduly large influence over the estimates. Since it is the genes with a low to moderate number of observations that benefit from shrinking, some modest bias of this sort is actually desired – but not as strong a bias as *w*(*n*) = 1 exhibits, and one not so purely focused on genes with very few observations. We therefore use the weights 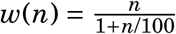, which initially increase linearly with the number of 1*+n*/100 observations, but eventually converge to 100 instead of increasing further. This has the desired effect of shifting the focus away from rarely observed genes, and concentrating it on genes with a moderate number of observations.

### Multiple initial copies

If of each distinct molecules the sample initially contains *R* > 1 identical copies (e.g. *R* = 2 if the initial molecules are double-stranded), each of these copies can be imagined to be amplified by a separate and independent PCR processes. But since the molecules are indistinguishable, these processes cannot be observed individually – we can observe only the (re-normalized) sum of the resulting family sizes. These observed normalized family size distribution is thus the average of *R* independent versions of *F*, and its variance is thus one *R*-th of the variance in equation 6, i.e.

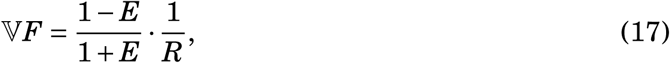

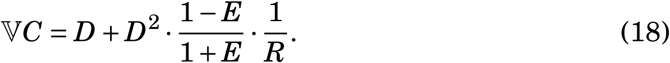

The density of distribution of *F* for *R* > 1 is the *R*-fold self-convolution of the density of *F* with itself (re-scaled to again have expected value one), and can thus be computed from the pre-computed matrix for the single-molecule case without performing additional simulations.

Parameter estimation proceeds just as for *R* = 1, except that when computing estimate *v′* of *VF*, we must now account for the reduction of the observed variance of *F* by a factor of 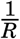 and equation 12 is thus replaced by

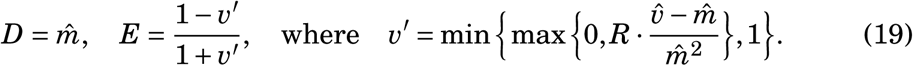

### Data Analysis

The reads from each of the downloaded sequenced libraries, were mapped (ignoring the barcode part) with *NGM* v0.5.2^13^ to the reference transcriptome of *D. melanogaster* (R6.08) respectively *E. coli* (strain K-12 MG1655). To avoid ambiguities during mapping for genes with multiple isoforms, we filtered the *D. melanogaster* transcriptome to contain only a single transcript per gene before mapping. For each gene, we picked either the single transcript with a FlyBase score of at least “moderately supported”, or the longest transcript (if multiple ones had score “moderately supported” or higher). After mapping the reads, we used the combination of mapping coordinates (both start and end for the paired-end *E. coli* data, only start for the single-end *D. melanogaster* data) and barcode (on both ends in the case of *E. coli*) as UMI. To account for sequencing errors, we merged similar UMIs (barcodes differing at most in one position, mapping coordinates by at most 30 bases for paired-end, 5 for sing-end libraries) using the graph-based algorithm of Smith *et al.*^9^. For the *E. coli* data we additionally combined reciprocal UMIs stemming from the two strands of a single template molecule, but stored the read counts for plus- and minus-strand separately (see Shiroguchi *et al.*^8^).

This yielded, for each of the libraries, a table comprising the gene id, startend end position, barcode and read-count(s) of each detected UMI. Based on this table, the error-correction thresholds (*T* = 5 for *E. coli*, *T* = 5 for *D. melanogaster* R1, *T* = 2 for *D. melanogaster* R2), and the initial number of molecules (actually, strands) for each UMI (*R* = 1 for *E. coli* due to the Y-shaped adapters, *R* = 2 for D. melanogaster due to secondary strand synthesis before amplification) our algorithm computed library-wide and raw as well as shrunken gene-specific estimates of the reaction efficiency, of the average number of reads per UMI, and of the loss. For the *E. coli* data, the error-correction threshold was applied to the plus- and minus-strand read counts separately, filtering out UMIs if either count lay below the chosen threshold. This increased the loss of true UMIs, and we modified the definition of the loss accordingly to 𝓁 = 1 − (1 − ℙ(*C < T*))^2^ (compare to equation 10). In addition to the gene-specific parameter and loss estimates, our algorithm output the observed number of UMIs 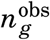 and the estimated total number of UMIs (i.e. transcript molecules) 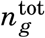.

### Simulation

We determined the residual error of the corrected transcript counts using a simulation approach. We started from the (loss-corrected) estimated transcript counts 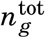 and (shrunken) gene-specific estimates for reaction efficiency *Ê_g_* and sequencing depth 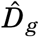*_g_* of gene *g* ∈ {1,…, *K*} that we computed for replicate 1 of the *D. melanogaster* dataset. First we rounded *n*^tot^ to the next number in the series 10, 30, 100, 300,… and used the resulting number as the *true* number 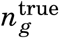 of transcripts of gene *g*. For each gene *g*, we then used the amplification+sequencing model (with parameters *E_g_*, *D_g_* and *R* = 2 meaning double-stranded molecules) to simulate the sequencing of 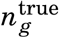 UMIs, which yielded for each gene 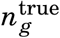 read counts, one for each UMI. To this list comprising gene id and (for each gene) 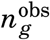 read counts, we applied our algorithm, using *T* = 5 and *R* = 2 as before (but passing along no other information from the first run of the algorithm). The algorithm thus dropped all UMIs with fewer than *T* = 5 reads, treated the remaining UMIs for each gene *g* as the *observed* number of UMIs 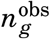, re-estimated the (shrunken) gene-specific losses, and used them to correct 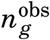 for these losses to arrive at an estimated total transcript count 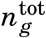. Finally, we computed for each gene the *relative quantification error* as

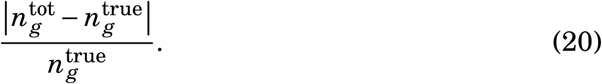

## SUPPLEMENTAL FIGURES

**Figure S1:**
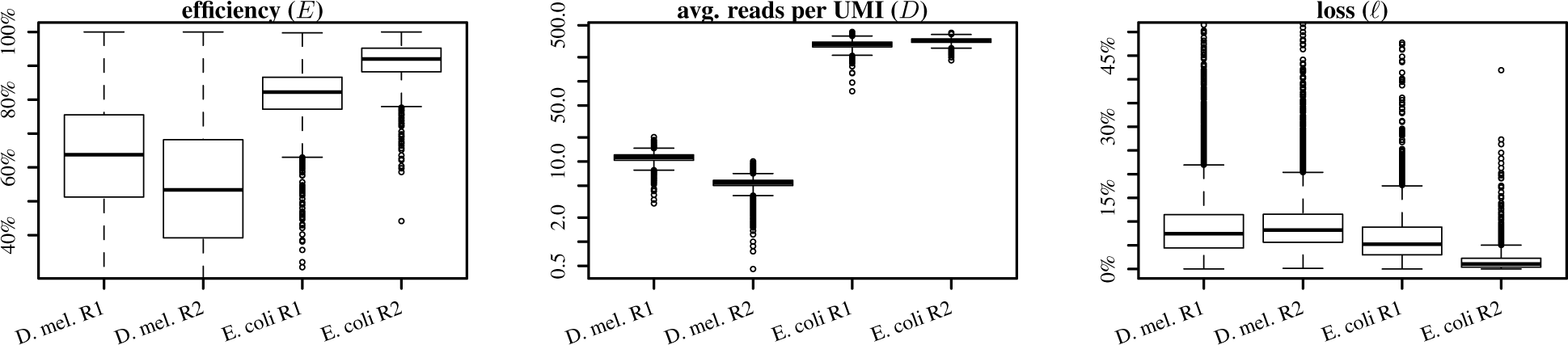
Variability of the raw (unshrunken) model parameters and resulting loss between genes. Includes parameter for 7481 detected genes in *D. mel.* R1, 8001 genes in R2, 2380 genes in *E. coli* R1 and 2308 genes in R2.

**Figure S2:**
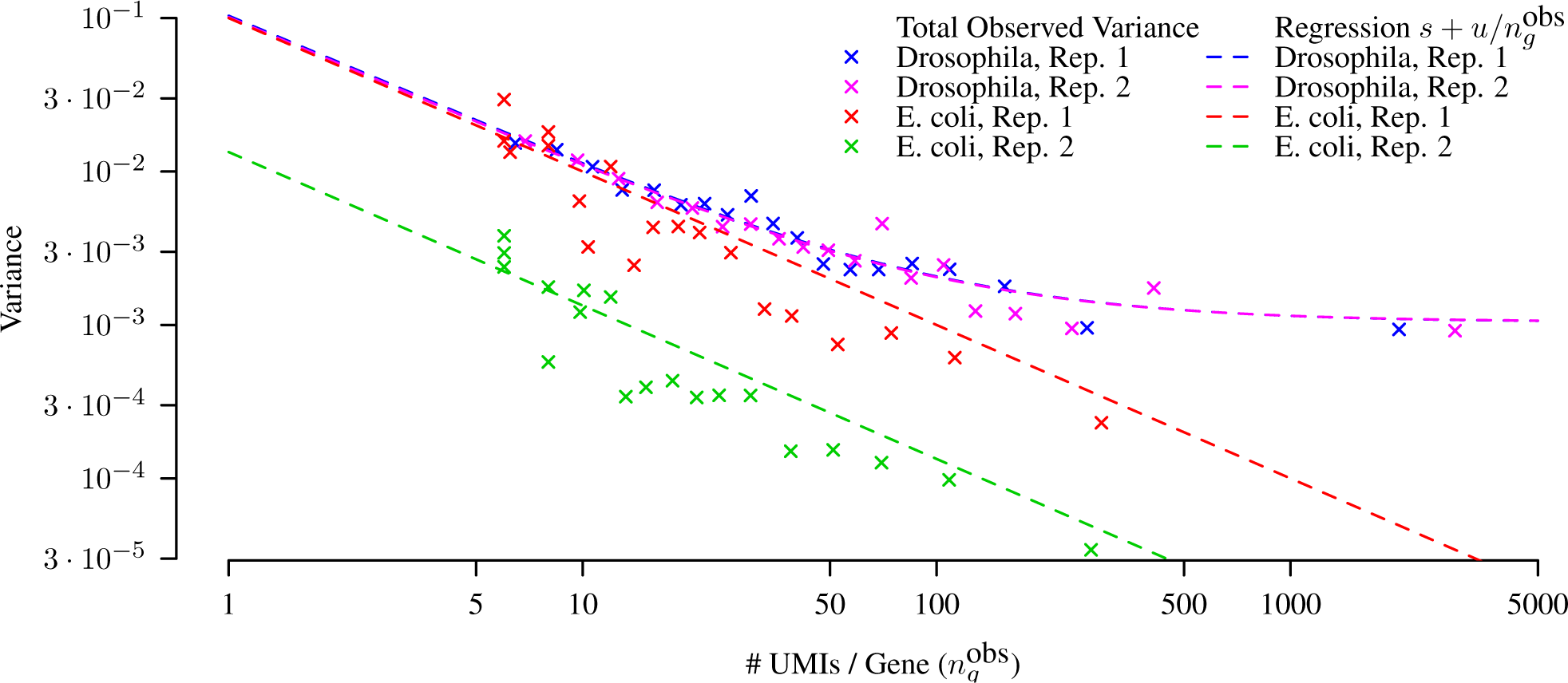
Total variance of the raw gene-specific loss estimates. Total observed variance was computed for bins containing 20 genes with a similar number 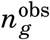 of observed true UMIs. The regression curve *s + u*/*n*^obs^ used to infer the optimal gene-specific shrinkage factors *λ_g_* comprises two components, the variance *s* of the loss between genes, and the 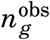-dependent error of the (raw) gene-specific loss estimates 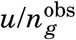. See also *Gene-specific estimates & corrections*.

**Figure S3:**
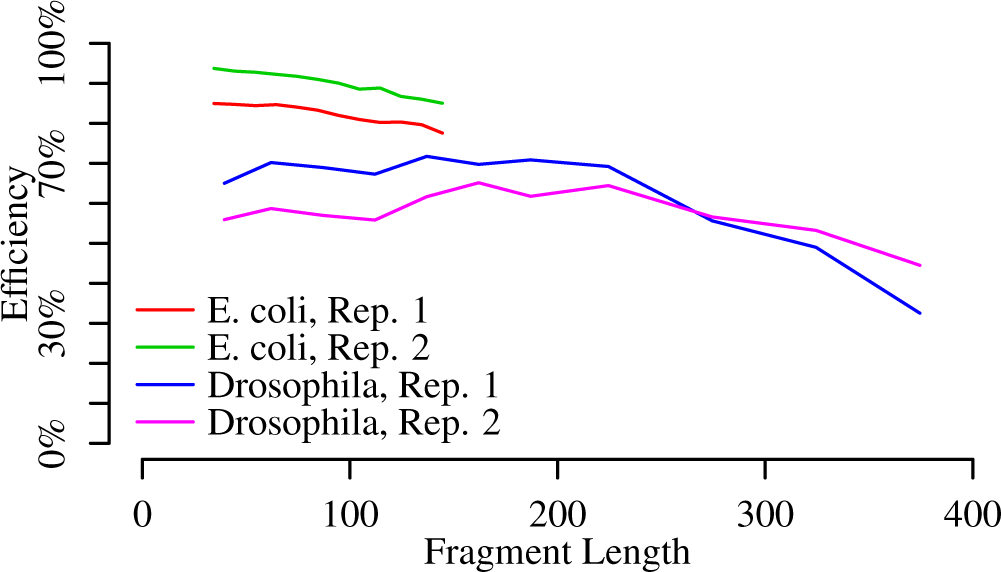
Length dependence of PCR efficiency. For each experiment, the detected UMIs were binned according to fragment length, and the PCR efficiency estimated independently for each bin.

